# A simple representation of three-dimensional molecular structure

**DOI:** 10.1101/136705

**Authors:** Seth D. Axen, Xi-Ping Huang, Elena L. Cáceres, Leo Gendelev, Bryan L. Roth, Michael J. Keiser

## Abstract

Statistical and machine learning approaches predict drug-to-target relationships from 2D small-molecule topology patterns. One might expect 3D information to improve these calculations. Here we apply the logic of the Extended Connectivity FingerPrint (ECFP) to develop a rapid, alignment-invariant 3D representation of molecular conformers, the Extended Three-Dimensional FingerPrint (E3FP). By integrating E3FP with the Similarity Ensemble Approach (SEA), we achieve higher precision-recall performance relative to SEA with ECFP on ChEMBL20, and equivalent receiver operating characteristic performance. We identify classes of molecules for which E3FP is a better predictor of similarity in bioactivity than is ECFP. Finally, we report novel drug-to-target binding predictions inaccessible by 2D fingerprints and confirm three of them experimentally with ligand efficiencies from 0.442 - 0.637 kcal/mol/heavy atom.

## Introduction

Many molecular representations have arisen since the early chemical informatics models of the 1970s, yet the most widely used still operate on the simple two-dimensional (topological) structures of small molecules. Fingerprints, which encode molecular 2D substructures as overlapping lists of patterns, were a first means to scan chemical databases for structural similarity using rapid bitwise logic on pairs of molecules. Pairs of molecules that are structurally similar, in turn, often share bioactivity properties ^1^ such as protein binding profiles. Whereas the prediction of biological targets for small molecules would seem to benefit from a more thorough treatment of a molecule’s explicit ensemble of three-dimensional (3D) conformations ^2^, pragmatic considerations such as calculation cost, alignment invariance, and uncertainty in conformer prediction ^3^ nonetheless limit the use of 3D representations by large-scale similarity methods such as the Similarity Ensemble Approach (SEA) ^4,5^, wherein the count of pairwise molecular calculations reaches into the hundreds of billions. Furthermore, although 3D representations might be expected to outperform 2D ones, in practice, 2D representations nonetheless are in wider use and can match or outperform them ^3,6–8^.

The success of statistical and machine learning approaches building on 2D fingerprints reinforces the trend. Naive Bayes Classifiers (NB) ^9–11^, Random Forests (RF) ^12,13^, Support Vector Machines (SVM) ^10,14,15^, and Deep Neural Networks (DNN) ^16–20^ predict a molecule’s target binding profile and other properties from the features encoded into its 2D fingerprint. SEA and methods building on it such as Optimized Cross Reactivity Estimation (OCEAN) ^21^ quantify and statistically aggregate patterns of molecular pairwise similarity to the same ends. Yet these approaches cannot readily be applied to the 3D molecular representations most commonly used. The Rapid Overlay of Chemical Structures (ROCS) method is an alternative to fingerprints that instead represents molecular shape on a conformer-by-conformer basis via gaussian functions centered on each atom. These functions may then be compared between a pair of conformers ^22,23^. ROCS however must align conformers to determine pairwise similarity; in addition to the computational cost of each alignment, which linear algebraic approximations such as SCISSORS ^24^ mitigate, the method provides no invariant fixed-length fingerprint (feature vectors) per molecule or per conformer for use in machine learning. One way around this limitation is to calculate an all-by-all conformer similarity matrix ahead of time, but this is untenable for large datasets such as ChEMBL ^25^ or the 70-million datapoint ExCAPE-DB ^26^, especially as the datasets continue to grow.

Feature Point Pharmacophores (FEPOPS), on the other hand, use *k*-means clustering to build a fuzzy representation of a conformer using a small number of clustered atomic feature points, which simplify shape and enable rapid comparison ^27,28^. FEPOPS excels at scaffold hopping, and it can use charge distribution based pre-alignment to circumvent a pairwise alignment step. However, pre-alignment can introduce similarity artifacts, such that explicit pairwise shape-based or feature-point-based alignment may nonetheless be preferred ^27^. Accordingly, 3D molecular representations and scoring methods typically align conformers on a pairwise basis ^2,3^. An alternative approach is to encode conformers against 3- or 4-point pharmacophore keys that express up to 890,000 or 350 million discrete pharmacophores, respectively ^29,30^. The count of purchasable molecules alone, much less their conformers, however, exceeds 200 million in databases such as ZINC (zinc.docking.org) ^31^, and the structural differences determining bioactivity may be subtle.

To directly integrate 3D molecular representations with statistical and machine learning methods, we developed a 3D fingerprint that retains the advantages of 2D topological fingerprints. Inspired by the widely used circular ECFP (2D) fingerprint, we develop a spherical Extended 3D FingerPrint (E3FP) and assess its performance relative to ECFP for various systems pharmacology tasks. E3FP is an open-source fingerprint that encodes 3D information without the need for molecular alignment, scales linearly with 2D fingerprint pairwise comparisons in computation time, and is compatible with statistical and machine learning approaches that have already been developed for 2D fingerprints. We use it to elucidate regions of molecular similarity space that could not previously be explored. To demonstrate its utility, we combine E3FP with SEA to predict novel target-drug activities that SEA could not discover using ECFP, and confirm experimentally that they are correct.

## Results

The three-dimensional fingerprints we present are motivated by the widely-used two-dimensional (2D) Extended Connectivity FingerPrint (ECFP) ^32^, which is based on the Morgan algorithm ^33^. ECFP is considered a 2D or “topological” approach because it encodes the internal graph connectivity of a molecule without explicitly accounting for 3D structural patterns the molecule may adopt in solution or during protein binding. While ECFP thus derives from the neighborhoods of atoms directly connected to each atom, a 3D fingerprint could incorporate neighborhoods of nearby atoms in 3D space, even if they are not directly bonded. We develop such an approach and call it an Extended Three-Dimensional FingerPrint (E3FP).

## A single small molecule yields multiple 3D fingerprints

Many small molecules can adopt a number of energetically favorable 3D conformations, termed “conformers”. In the absence of solved structures, it is not always apparent which conformer a molecule will adopt in solution, how this may change on protein binding, and which protein-ligand interactions may favor which conformers ^34^. Accordingly, we generate separate E3FPs for each of multiple potential conformers per molecule. E3FP encodes all three-dimensional substructures from a single conformer into a bit vector, represented as a fixed-length sequence of 1s and 0s (Figure 1a). This is analogous to the means by which ECFP represent two-dimensional substructures. To encode the three-dimensional environment of an atom, E3FP considers information pertaining not only to contiguously bound atoms, but also to nearby unbound atoms and to relative atom orientations (stereochemistry). We designed this process to be minimally sensitive to minor structural fluctuations, so that conformers could be distinguished while the set of conformers for a given molecule would retain a degree of internal similarity in E3FP space.

**Figure 1.**
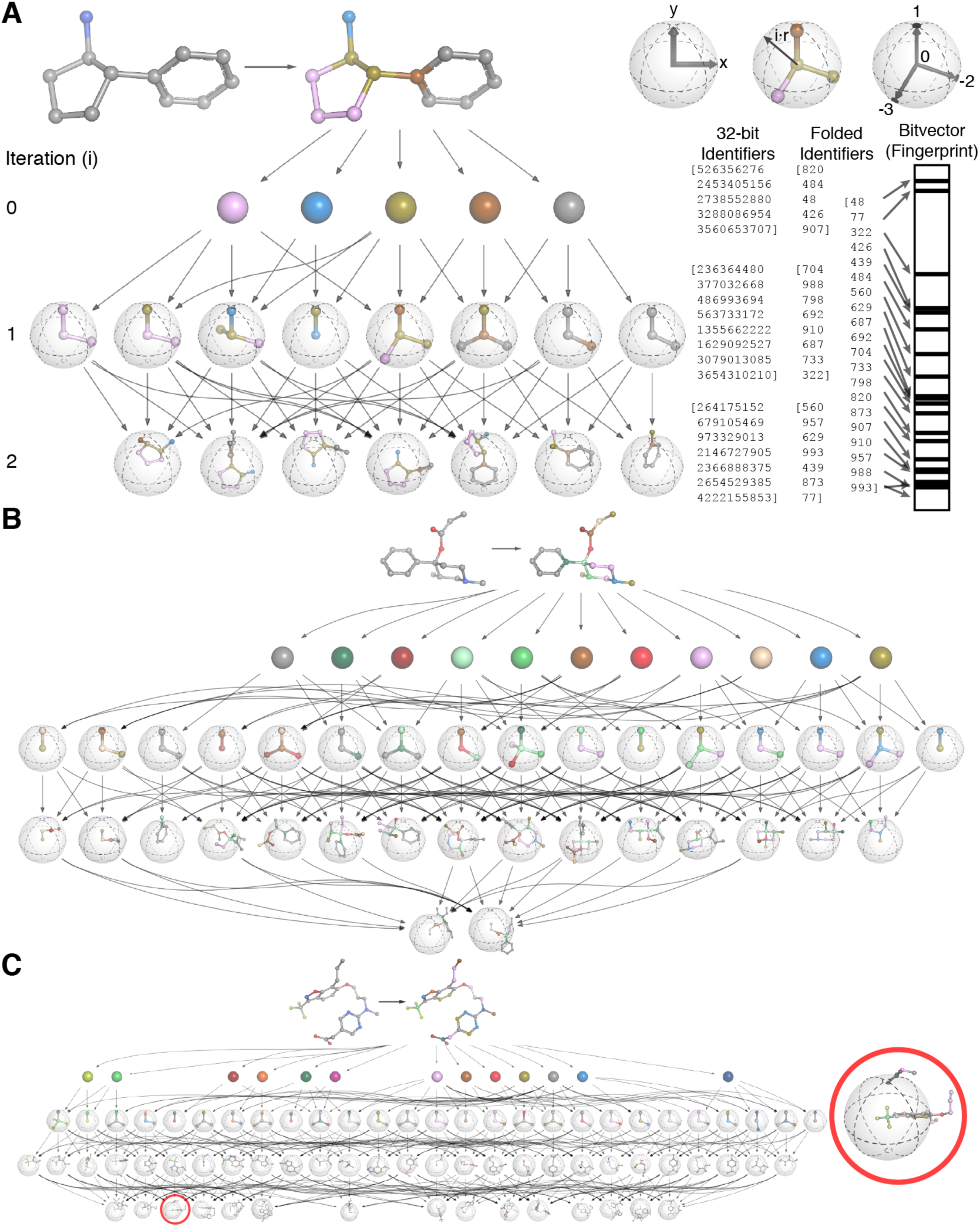
Diagram of information flow in the E3FP algorithm. A) Overview of fingerprinting process for cypenamine. At iteration 0, we assign atom identifiers using a list of atomic invariants and hash these into integers (shown here also as unique atom colors). At iteration *i*, shells of radius *i · r* center on each atom (top right). The shell contains bound and unbound neighbor atoms. Where possible, we uniquely align neighbor atoms to the *xy*-plane (top right) and assign stereochemical identifiers. Convergence occurs when a shell’s substructure contains the entire molecule (third from the right) or at the maximum iteration count. Finally we “fold” each iteration’s substructure identifiers to 1024-bit space. B) Overview of fingerprinting for compound **1**. C) Overview of fingerprinting for a large, flexible molecule (CHEMBL210990; expanded in Figure S1). A three-dimensional substructure can consist of two disconnected substructures and their relative orientations (right).

The binding-relevant conformers of most small molecules are not known *a priori*. Accordingly, prior to constructing any 3D fingerprint, we generate a library of potential conformers for the molecule, each of which in turn will have a unique fingerprint. We employed a previously published protocol using the open-source RDKit package ^35^, wherein the authors determined the number of conformers needed to recover the correct ligand conformation from a crystal structure as a function of the number of rotatable bonds in the molecule, with some tuning (see Experimental Section).

## E3FP encodes small molecule 3D substructures

The core intuition of E3FP generation (Figure 1a) is to draw concentrically larger shells and encode the 3D atom neighborhood patterns within each of them. To do so, the algorithm proceeds from small to larger shells iteratively. First, as in ECFP, we uniquely represent each type of atom and the most important properties of its immediate environment. To do so, we assign 32-bit integer identifiers to each atom unique to its count of heavy atom immediate neighbors, its valence minus neighboring hydrogens, its atomic number, its atomic mass, its atomic charge, its number of bound hydrogens, and whether it is in a ring. This can result in many fine-grained identifiers, some examples of which are visualized as differently colored atoms for the molecule cypenamine in Figure 1a and for larger molecules in Figure 1b-c.

At each subsequent iteration, we draw a shell of increasing radius around each atom, defining the neighbors as the atoms within the shell as described above. The orientation and connectivity of the neighbors--or lack thereof (as in Figure 1c, red circle, expanded in Figure S1)--is combined with the neighbors’ own identifiers from the previous iteration to generate a new joint identifier. Thus, at any given iteration, the information contained within the shell is the union of the substructures around the neighbors from the previous iterations merged with the neighbors’ orientation and connectivity with respect to the center atom of the current shell. The set of atoms represented by an identifier therefore comprise a three-dimensional substructure of the molecule.

We continue this process up to a predefined maximum number of iterations or until we have encountered all substructures possible within that molecule. We then represent each identifier as an “on” bit in a sparse bit vector representation of the entire conformer (Figure 1a, bitvector). Each “on” bit indicates the presence of a specific three-dimensional substructure. The choice of numerical integer to represent any identifier is the result of a hash function (see Experimental Section) that spreads the identifiers evenly over a large integer space. Because there are over four billion possible 32-bit integers and we observe far fewer than this number of molecular substructures (identifiers) in practice, each identifier is unlikely to collide with another and may be considered unique to a single atom or substructure. Since this still remains a mostly empty identifier space, we follow the commonly used approach from ECFP, and “fold” E3FP down to a shorter bitvector for efficient storage and swift comparison; adapting the 1024-bit length that has been effective for ECFP4 ^6,36^ (Table S2).

To demonstrate the fingerprinting process, Figure 1a steps through the generation of an E3FP for the small molecule cypenamine. First, four carbon atom types and one nitrogen atom type are identified, represented by five colors. As cypenamine is fairly small, E3FP fingerprinting terminates after two iterations, at which point one of the substructures consists of the entire molecule. The slightly larger molecule **1** (CHEMBL270807) takes an additional iteration to reach termination (Figure 1b). Figure 1c and Figure S1 demonstrate the same process for 2-[2-[methyl-[3-[[7-propyl-3-(trifluoromethyl)-1,2-benzoxazol-6-yl]oxy]propyl]amino]pyrimidin-5-yl]acetic acid (CHEMBL210990). This molecule is more complex, with 13 distinct atom types, and in the conformation shown reaches convergence in three iterations. Because the molecule bends back on itself, in the second and third iterations, several of the identifiers represent substructures that are nearby each other in physical space but are not directly bound to each and other and indeed are separated by many bonds (e.g., red circle in Figure 1c). 2D fingerprints such as ECFP are inherently unaware of unconnected proximity-based substructures, but they are encoded in E3FP.

## SEA 3D fingerprint performance exceeds that of 2D in binding prediction

We were curious to determine how molecular similarity calculations using the new E3FP representations would compare to those using the 2D but otherwise similarly-motivated ECFP4 fingerprints. Specifically, we investigated whether the 3D fingerprint encoded information that would enhance performance over its 2D counterpart in common chemical informatics tasks.

The ECFP approach uses several parameters, (e.g., ECFP4 uses a radius of 2), and prior studies have explored their optimization ^36^. We likewise sought appropriate parameter choices for E3FP. In addition to the conformer generation choices described above, E3FP itself has four tunable parameters: 1) a shell radius multiplier (*r* in Figure 1a), 2) number of iterations (*i* in Figure 1a), 3) inclusion of stereochemical information, and 4) final bitvector length (1024 in Figure 1a). We explored which combinations of conformer generation and E3FP parameters produced the most effective 3D fingerprints for the task of recovering correct ligand binders for over 2,000 protein targets using the Similarity Ensemble Approach (SEA). SEA compares sets of fingerprints against each other using Tanimoto coefficients (TC) and determines a *p-value* for the similarity among the two sets; it has been used to predict drug off-targets ^4,5,37,38^, small molecule mechanisms of action ^39–41^, and adverse drug reactions ^4,42,43^. For the training library, we assembled a dataset of small molecule ligands that bind to at least one of the targets from the ChEMBL database with an *IC*_50_ of 10 μM or better. We then generated and fingerprinted the conformers using each E3FP parameter choice, resulting in a set of conformer fingerprints for each molecule and for each target. We performed a stratified 5-fold cross-validation on a target-by-target basis by setting aside one fifth of the known binders from a target for testing, searching this one fifth (positive data) and the remaining non-binders (negative data) against the target using SEA, and then computing true and false positive rates at all possible SEA *p-value* cutoffs. For each target in each fold, we computed the precision recall curve (PRC), the receiver operating characteristic (ROC), and the area under each curve (AUC). Likewise, we combined the predictions across all targets in a cross-validation fold to generate fold PRC and ROC curves.

As there are far more negative target-molecule pairs in the test sets than positives, a good ROC curve was readily achieved, as many false positives must be generated to produce a high false positive rate. Conversely, in such a case, the precision would be very low. We therefore expected the AUC of the PRC (AUPRC) to be a better assessment of parameter set ^44^. To simultaneously optimize for both a high AUPRC and a high AUC of the ROC (AUROC), we used the sum of these two values as the objective function, AUC_SUM_. We employed the Bayesian optimization program Spearmint ^45^ to optimize four of five possible E3FP parameters (we did not optimize fingerprint bit length, for simplicity of comparison to ECFP fingerprints) so as to maximize the AUC_SUM_ value and minimize runtime of fingerprinting (Figure S2).

We constrained all optimization solely to the choice of fingerprint parameters, on the same underlying collection of precomputed molecular conformers. For computational efficiency, we split the optimization protocol into two stages (see Experimental Section). This yielded an E3FP parameter set that used the three lowest energy conformers, a shell radius multiplier of 1.718, and 5 iterations of fingerprinting (Figure S4). After bootstrapping with 5 independent repeats of 5-fold cross-validation using E3FP, and ECFP4 on a larger set of 308,316 ligands from ChEMBL20, E3FP produced a mean AUPRC of 0.6426, exceeding ECFP4’s mean AUPRC of 0.5799 in the same task (Figure 2c,e; Table 1). Additionally, E3FP’s mean AUROC of 0.9886 exceeds ECFP4’s AUPRC of 0.9882 (Figure 2d-e; Table 1). Thus, at a SEA *p-value* threshold *p* ≤ 3.45×10^-47^, E3FP achieves an average *sensitivity* of 0.6976, *specificity* of 0.9974, *precision* of 0.5824, and *F*_*1*_ score of 0.6348. ECFP4 achieves 0.4647, 0.9986, 0.6236, and 0.5325, at this *p-value* threshold. ECFP4 is unable to achieve the high *F*_*1*_ score of E3FP, but at its maximum *F*_*1*_ score of 0.5896 it achieves a *sensitivity* of 0.6930, a specificity of 0.9966, and a *precision* of 0.5131 using a *p-value* threshold *p* ≤ 3.33×10^-23^. To ensure a fair comparison, we subjected ECFP to a grid search on its radius parameter and found that no radius value outperforms ECFP4 with both AUPRC and AUROC (Table S1). Additionally, fingerprints with longer bit lengths did not yield significant performance increases for E3FP or ECFP4, despite the expectation that longer lengths would lower feature collision rates (Table S2); indeed, it appears that increasing the fingerprint length reduced the performance of E3FP. By design, this optimization and consequent performance analysis does not attempt to quantify novelty of the predictions, nor assess the false negative or untested-yet-true-positive rate of either method.

**Figure 2.**
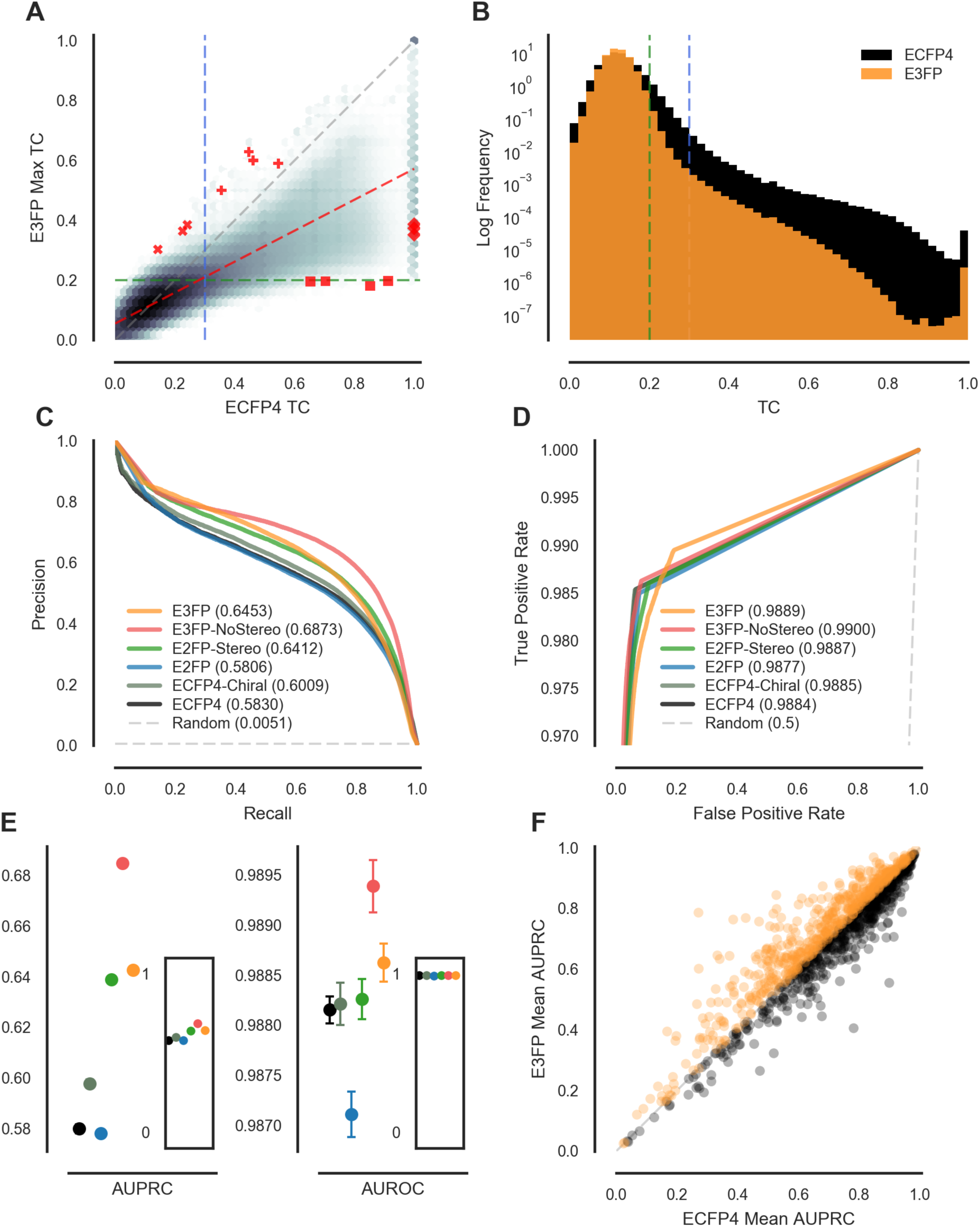
Comparative performance of E3FP and ECFP. For all pairs of 308,315 molecules from ChEMBL20, A) log density plot summarizing 95 billion maximum Tanimoto Coefficients (TC) calculated between E3FP conformer fingerprint sets versus corresponding TC by ECFP4 fingerprints. The dotted red line is a linear least squares fit. Optimal SEA TC cutoffs for E3FP (green) and ECFP4 (blue) are dotted lines. Red markers indicate examples in Figure 3. B) Histograms of TCs from (A). C) Combined precision-recall (PRC) curves from 5 independent 5-fold cross-validation runs using 1024-bit E3FP, E3FP without stereochemical identifiers (E3FP-NoStereo), E3FP without stereochemical identifiers or nearby unbound atoms (E2FP), E3FP without nearby unbound atoms (E2FP-Stereo), ECFP4, and ECFP4 with distinct bond types encoding chirality (ECFP4-Chiral). Only the PRC of the highest AUC fold is shown. D) Combined highest-AUC ROC curves for the same sets as in (C). E) Results of bootstrapping AUCs as in Table 1. Dots indicate mean AUC, and whiskers standard deviations. Insets show absolute scale. F) Target-wise comparison of mean AUPRCs using E3FP versus ECFP4.

**Table 1.** Performance of variations of E3FP and ECFP using SEA.

| Name | Mean Fold AUPRC | Mean Fold AUROC | Mean Target AUPRC | Mean Target AUROC |
| --- | --- | --- | --- | --- |
| ECFP4 | 0.5799 ± 0.0018 | 0.9882 ± 0.0001 | 0.6965 ± 0.2099 | 0.9772 ± 0.0387 |
| ECFP4-Chiral | 0.5977 ± 0.0017 | 0.9882 ± 0.0002 | 0.7021 ± 0.2088 | 0.9769 ± 0.0391 |
| E2FP | 0.5781 ± 0.0015 | 0.9871 ± 0.0002 | 0.7080 ± 0.2034 | 0.9768 ± 0.0392 |
| E2FP-Stereo | 0.6390 ± 0.0011 | 0.9883 ± 0.0001 | 0.7140 ± 0.2016 | 0.9780 ± 0.0371 |
| E3FP-NoStereo | 0.6849 ± 0.0012 | 0.9894 ± 0.0003 | 0.7312 ± 0.1989 | 0.9774 ± 0.0409 |
| E3FP | 0.6426 ± 0.0016 | 0.9886 ± 0.0002 | 0.7046 ± 0.1991 | 0.9805 ± 0.0326 |

Mean and standard deviations for combined fold AUPRC and AUROC curves versus target-wise AUPRC and AUROC curves across 5 independent repeats of 5-fold cross-validation are shown. A random classifier will produce a mean AUPRC of 0.0051 (fraction of positive target/mol pairs in test data), a mean target AUPRC of 0.0053 ± 0.0076, and a mean AUROC and mean target-wise AUROC of 0.5.

We note that E3FP was optimized here for use with SEA, and SEA inherently operates on sets of fingerprints, such as those produced when fingerprinting a set of conformers. Most machine learning methods, however, operate on individual fingerprints. To determine how well E3FP could be integrated into this scenario, we repeated the entire cross-validation with four common machine learning classifiers: Naive Bayes Classifiers (NB), Random Forests (RF), Support Vector Machines with a linear kernel (LinSVM), and Artificial Neural Networks (NN). As these methods process each conformer independently, we computed the maximum score across all conformer-specific fingerprints for a given molecule, and used that score for cross-validation. Compared to cross-validation with SEA, LinSVM and RF produced better performance by PRC using both E3FP and ECFP4, while NB and RF suffered a performance loss (Figure S5). For ECFP4, this trend continued when comparing ROC curves, while for E3FP it did not (Figure S6). In general, the machine learning methods underperformed when using E3FP compared to ECFP4. When we instead took the bitwise mean of all conformer-specific E3FPs to produce one single summarizing “float” fingerprint per molecule, we observed an improvement across all machine learning methods except for LinSVM. The most striking difference was for RF, where performance with “mean E3FP” then matched ECFP4.

## 3D fingerprints encode different information than their 2D counterparts

2D fingerprints such as ECFP4 may denote stereoatoms using special disambiguation flags or identifiers from marked stereochemistry (here termed “ECFP4-Chiral”) ^32^. E3FP encodes stereochemistry more natively. Conceptually, all atoms within a spatial “neighborhood” and their relative orientations within that region of space are explicitly considered when constructing the fingerprint. To quantify how stereochemical information contributes to E3FP’s improved AUPRC over that of ECFP4, we constructed three “2D-like” limited variants of E3FP, each of which omits some 3D information and is thus more analogous to ECFP. The first variant, which we term “E2FP,” is a direct analogue of ECFP, in which only information from directly bound atoms are included in the identifier and stereochemistry is ignored. This variant produces similar ROC and PRC curves to that of ECFP4 (Figure 2c-d; Figures S7-S8). A second variant, “E2FP-Stereo,” includes information regarding the relative orientations of bound atoms. E2FP-Stereo achieves a performance between that of ECFP4 and E3FP, demonstrating that E3FP’s approach for encoding stereochemical information is effective (Figure 2c-d). The third variant, “E3FP-NoStereo,” includes only the information from bound and unbound atoms. E3FP-NoStereo performs slightly better than E3FP in both ROC and PRC analysis (Figure 2c-d), indicating that E3FP’s enhanced performance over ECFP4 in PRC analysis is due not only to the relative orientations of atoms but also due to the inclusion of unbound atoms. All variants of E3FP with some form of 3D information outperformed both ECFP4 and ECFP4-Chiral (Figure 2c-d; Figures S7-S8).

On average, the final E3FP parameters yield fingerprints with 35% more “on” bits than ECFP4, although if run for the same number of iterations, ECFP is denser. Thus E3FP typically runs for more iterations (Figure S4c-d). Folding E3FP down to 1024 bits results in an average loss of only 1.4 bits to collisions. The TCs for randomly chosen pairs of molecules by E3FP are generally lower (Figure 2a-b) than those for ECFP4, and there are fewer molecules with identical fingerprints by E3FP than by ECFP4. The final E3FP parameter set outperforms ECFP up to the same number of iterations (Table S1, Figure 2c-d). Intriguingly, E3FP outperforms ECFP4 at this task on a per-target basis for a majority of targets (Figure 2f).

## Fourteen molecular pairs where 3D and 2D fingerprints disagree

To explore cases where E3FP and ECFP4 diverge, we computed E3FP versus ECFP4 pairwise similarity scores (Tanimoto coefficients; TCs) for all molecule pairs in ChEMBL20 (red markers in Figure 2a). We then manually inspected pairs from four regions of interest. Pairs representative of overall trends were selected, with preference toward pairs that had been assayed against the same target (Table S3). The first region contains molecule pairs with TCs slightly above the SEA significance threshold for E3FP but below the threshold for ECFP4 (denoted by ‘x’ markers). These predominantly comprise small compact molecules, with common atom types across multiple orders or substituents on rings (Figure 3a). Some of these molecules are already reported to interact with the same protein targets. For instance, [(E)-3-aminoprop-1-enyl]phosphinic acid (CHEMBL113217) binds to GABA-B receptor with an *IC*_50_ of 280 nM, while 3-aminobutylphosphinic acid (CHEMBL113907) binds GABA-B with a similar *IC*_50_ of 500 nM (Figure 3a) ^46^. In another example, azocan-(2Z)-ylideneamine (CHEMBL329431) binds to inducible, brain, and endothelial human nitric-oxide synthases with *IC*_*50*_s of 10.0 μM, 10.1 μM, and 59 μM, respectively ^47^, while hexahydro-cyclopenta[c]pyrrol-(1Z)-ylideneamine (CHEMBL365849) binds to the same targets at 3.1 μM, 310 nM, and 4.7 μM ^48^. The black asterisk alongside this pair marks similar affinities for the first target (within 1 log), and the gold asterisks affinities for the second two, each spanning two logs. Red asterisks mark targets whose affinities differ by more than two logs, but no such cases were found for this region.

**Figure 3.**
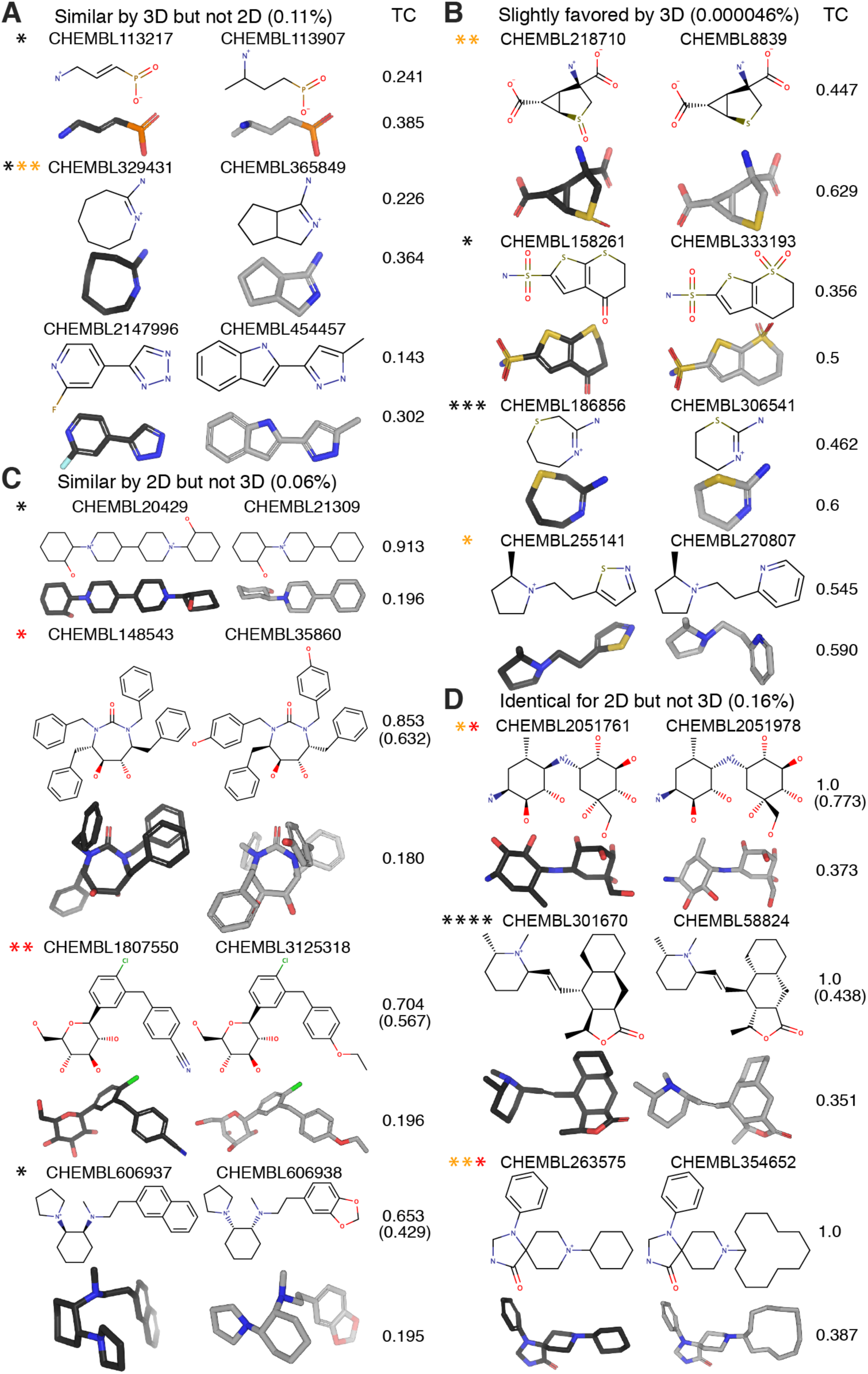
Examples of Molecule Pairs with High Differences between E3FP and ECFP Tanimoto Coefficients. Molecule pairs were manually selected from regions of interest, displayed as red markers in Figure 2a: A) Upper left, B) Upper right, C) Lower right, and D) Far right. Pair TCs for ECFP4 and E3FP are shown next to the corresponding 2D and 3D representations; the conformer pairs shown are those corresponding to the highest pairwise E3FP TC. Where pair TCs for ECFP4 with stereochemical information differ from standard ECFP4, they are included in parentheses. Each colored asterisk indicates a target for which existing affinity data for both molecules was found in the literature and is colored according to fold-difference in affinity: black for <10-fold, orange for 10-100-fold, red for >100-fold.

The second region (red crosses in Figure 2a) contains molecule pairs with TCs considered significant both in 2D and in 3D, but whose similarity was nonetheless greater by 3D (Figure 3b). For instance, the molecule pairs often differed by atom types in or substituents on a ring, despite a high degree of similarity in 3D structures. In the case of 4-oxo-5,6-dihydrothieno[2,3-b]thiopyran-2-sulfonamide (CHEMBL158261) and 7,7-dioxo-5,6-dihydro-4H-thieno[2,3-b]thiopyran-2-sulfonamide (CHEMBL333193), the molecules bind to carbonic anhydrase II with near-identical affinities of 3.6 nM and 3.3 nM ^49^. Interestingly, the 2D similarity of this pair is barely above the significance threshold. In another example, the molecules [1,4]thiazepan-(3E)-ylideneamine (CHEMBL186856) and [1,3]thiazinan-(2E)-ylideneamine (CHEMBL306541) achieve markedly similar pharmacological profiles, as the first binds to the inducible, brain, and endothelial human nitric-oxide synthases with *IC50*s of 1.2 μM, 2.8 μM, and 10.5 μM ^50^, whereas the second was reported at 2.9 μM, 3.2 μM, and 7.1 μM ^51^. On the other hand, two other pairs somewhat differ in binding profile: while 4-aminho-2-thiabicyclo(3.1.0)hexane-4,6-dicarboxylic acid (CHEMBL218710) binds to metabotropic glutamate receptors 2 and 3 with *K*_*i*_s of 508 nM and 447 nM, 2-thia-4-aminobicyclo(3.1.0)hexane-4,6-dicarboxylic acid (CHEMBL8839) binds to these targets more potently, at 40.6 nM and 4.7 nM ^52^. Likewise, the binding profiles of 5-[2-[(2R)-2-methylpyrrolidin-1-yl]ethyl]-1,2-thiazole (CHEMBL255141) and **1** to histamine H3 receptor differed by approximately an order of magnitude, with respective *K*_*i*_s of 17 nM and 200 nM ^53^.

The third region (red squares in Figure 2a) contains molecule pairs significant in 2D but not in 3D (Figure 3c), and the fourth region (red diamonds in Figure 2a) contains pairs identical by 2D yet dissimilar in 3D (Figure 3d). These examples span several categories: First, the conformer generation protocol failed for some pairs of identical or near-identical molecules having many rotatable bonds, because we generated an insufficient number of conformers to sample the conformer pair that would attain high 3D similarity between them (not shown). Second, in cases where the 2D molecules do not specify chirality, the specific force field used may favor different chiralities, producing artificially low 3D similarity. As an example, 2-[4-[1-(2-hydroxycyclohexyl)piperidin-4-yl]piperidin-1-yl]cyclohexan-1-ol (CHEMBL20429) and 2-(4-cyclohexylpiperidin-1-yl)cyclohexan-1-ol (CHEMBL21309) (Figure 3c) have relatively similar affinities for vesicular acetylcholine transporter at 200 nM and 40 nM ^54^ despite their low 3D similarity. Third, some pairs consist of molecules primarily differentiated by the size of one or more substituent rings (Figure 3c-d). ECFP4 is incapable of differentiating rings with 5 or more identical atom types and only one substituent, while E3FP substructures may include larger portions of the rings. The role of ring size is revealed in the target affinity differences for one such pair: 8-cyclohexyl-1-phenyl-1,3,8-triazaspiro[4.5]decan-4-one (CHEMBL263575) binds to the kappa opioid, mu opioid, and nociceptin receptors with *K*_*i*_s of 100 nM, 158 nM, and 25 nM, while 8-cyclododecyl-1-phenyl-1,3,8-triazaspiro[4.5]decan-4-one (CHEMBL354652) binds to the same receptors notably more potently at 2.9 nM, 0.28 nM, and 0.95 nM ^55^. Fourth, many pairs consist of molecules primarily differentiated by the order of substituents around one or more chiral centers (Figure 3c-d). The molecules (4S,5S,6S,7S)-1,3,4,7-tetrabenzyl-5,6-dihydroxy-1,3-diazepan-2-one (CHEMBL148543) and (4R,5S,6S,7R)-4,7-dibenzyl-5,6-dihydroxy-1,3-bis[(4-hydroxyphenyl)methyl]-1,3-diazepan-2-one (CHEMBL35860), for example, bind to HIV type 1 protease with disparate *Ki*s of 560 nM ^56^ and 0.12 nM ^57^ despite their exceptionally high 2D similarity of 0.853 TC. Likewise, 4-[[2-chloro-5-[(2S,3R,4R,5S,6R)-3,4,5-trihydroxy-6-(hydroxymethyl)oxan-2-yl]phenyl]methyl]benzonitrile (CHEMBL1807550) and (2S,3R,4S,5S,6R)-2-[4-chloro-3-[(4 ethoxyphenyl)methyl]phenyl]-2-deuterio-6-(hydroxymethyl)oxane-3,4,5-triol (CHEMBL3125318) have opposing specificities for the human sodium/glucose cotransporters 1 and 2; while the former has *IC*_*50*_s of 10 nM and 10 μM for the targets ^58^, the latter has *IC50*s of 3.1 μM and 2.9 nM ^59^. In another example, despite being identical by standard 2D fingerprints, the stereoisomers (1S,2S,3R,4S,5S)-5-[[(1R,2S,3S,4S,6S)-4-amino-2,3-dihydroxy-6-methylcyclohexyl]amino]-1-(hydroxymethyl)cyclohexane-1,2,3,4-tetrol (CHEMBL2051761) and (1S,2S,3R,4S,5S)-5-[[(1S,2S,3S,4S,6S)-4-amino-2,3-dihydroxy-6-methylcyclohexyl]amino]-1-(hydroxymethyl)cyclohexane1,2,3,4-tetrol (CHEMBL2051978) bind to maltase-glucoamylase with *IC*_*50*_s of 28 nM versus 1.5 μM, and to sucrase-isomaltase at 7.5 nM versus 5.3 μM ^60^. The stereoisomers (3S,3aS,4R,4aS,8aS,9aR)-4-[(E)-2-[(2R,6S)-1,6-dimethylpiperidin-2-yl]ethenyl]-3 methyl-3a,4,4a,5,6,7,8,8a,9,9a-decahydro-3H-benzo[f][2]benzofuran-1-one(CHEMBL301670) and (3S,3aR,4S,4aR,8aR,9aS)-4-[(Z)-2-[(2R,6S)-1,6-dimethylpiperidin-2-yl]ethenyl]-3-methyl-3a,4,4a,5,6,7,8,8a,9,9a-decahydro-3H-benzo[f][2]benzofuran-1-one (CHEMBL58824), however, show a case where 3D dissimilarity is a less effective guide, as both molecules bind to the muscarinic acetylcholine receptors M_1_-M_4_ with generally similar respective *IC*_*50*_s of 426.58 nM versus 851.14 nM, 95.5 nM versus 851.14 nM, 1.6 μM versus 794.33 nM, and 173.78 nM versus 794.33 nM ^61^. Similarly, (1S,2R)-N-methyl-N-(2-naphthalen-2-ylethyl)-2-pyrrolidin-1-ylcyclohexan-1-amine (CHEMBL606937) and (1R,2S)-N-[2-(1,3-benzodioxol-5-yl)ethyl]-N-methyl-2-pyrrolidin-1-ylcyclohexan-1-amine(CHEMBL606938) have low similarity in 3D but bind to sigma opioid receptor with *IC*_*50*_s of 37 and 34 nM ^62^.

## E3FP predicts correct new drug off-targets that are not apparent in 2D

As E3FP enhanced SEA performance in retrospective tests (Figure 2c-d), we hypothesized that this combination might identify novel interactions as yet overlooked with two-dimensional fingerprints. We therefore tested whether SEA with E3FP would make correct drug-to-target predictions that SEA with ECFP4 did not make. Using a preliminary choice of E3FP parameters (Table S4), we generated fingerprints for all in-stock compounds in the ChEMBL20 subset of the ZINC15 (zinc15.docking.org) database with a molecular weight under 800 Da. As our reference library, we extracted a subset of ChEMBL20 comprising 309 targets readily available for testing by radioligand binding assay in the Psychoactive Drug Screening Program (PDSP) ^63^ database. Using SEA on this library, we identified all drug-to-target predictions with a *p-value* stronger than 1×10^-25^. To focus on predictions specific to E3FP, we removed all predictions with a *p-value* stronger than 0.1 when counter-screened by SEA with ECFP4, resulting in 9,331 novel predicted interactions. We selected eight predictions for testing by binding assay; of these, five were inconclusive, and three bound to the predicted target subtype or to a close subtype of the same receptor (Table S4-S7). We address each of the latter in turn.

The E3FP SEA prediction that the psychostimulant and antidepressant ^64–66^ cypenamine (CHEMBL2110918, KEGG:D03629), for which we could find no accepted targets in the literature despite its development in the 1940s, would bind to the human nicotinic acetylcholine receptor (nAchR) α2β4 was borne out with a *K*_*i*_ of 4.65 μM (Figure 4c; Table S7). Of note, this corresponds to a high ligand efficiency (LE) of 0.610 kcal/mol/heavy atom (see Experimental Section). An LE greater than 0.3 kcal/mol/heavy atom is generally considered a promising drug candidate ^67^. As any prediction is only as specific as the reference ligand data from ChEMBL upon which it was based, we assayed cypenamine against multiple subtypes of nAchR. Cypenamine also bound to the nAchR subtypes α3β4 and α4β4 with *K*_*i*_’s of 2.69 and 4.11 μM (Figure 4d-e, Table S7) and ligand efficiencies of 0.637 and 0.616 kcal/mol/heavy atom.

**Figure 4.**
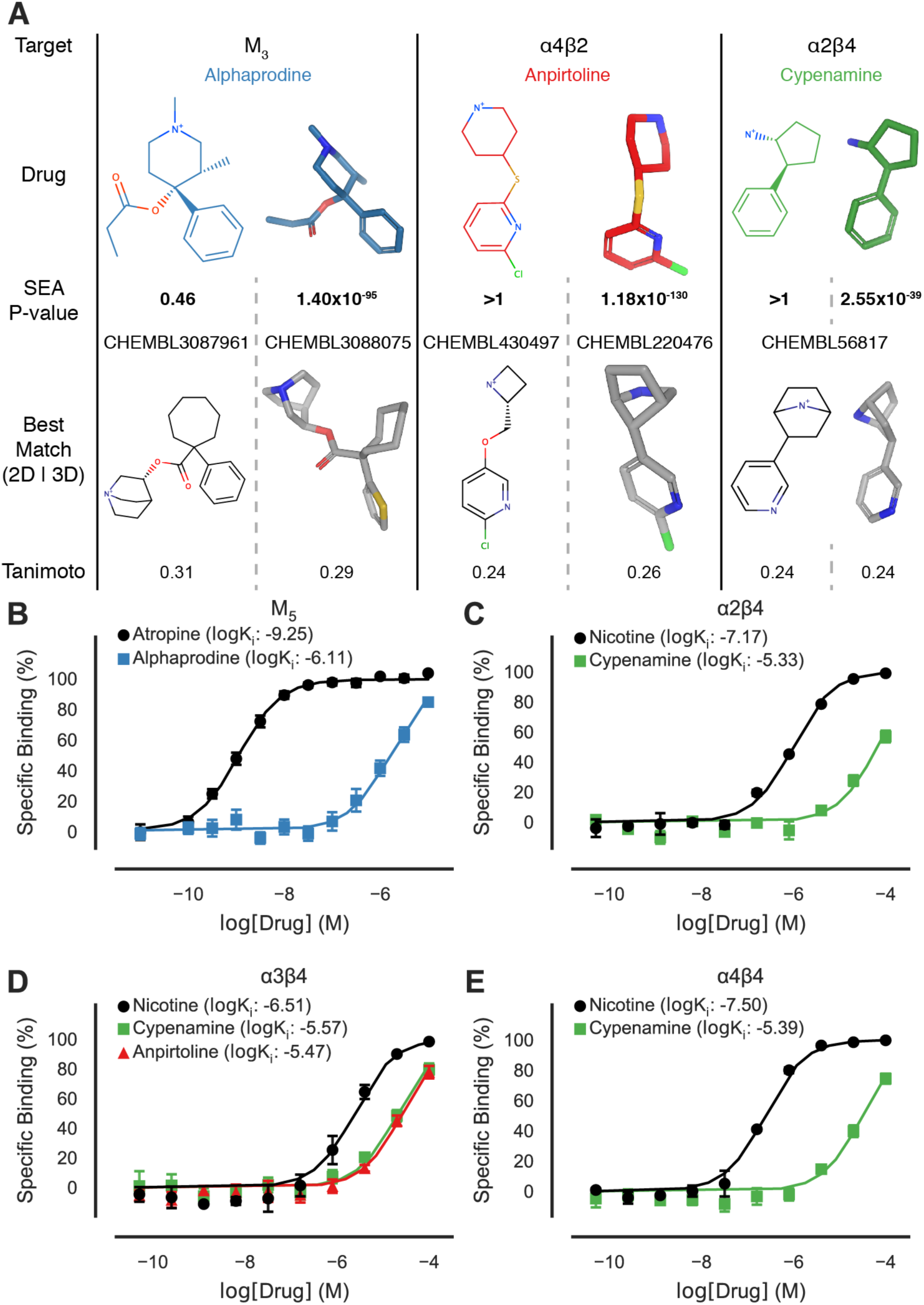
Experimental results of novel compound-target predictions. A) SEA predictions that motivated the binding experiments, with 2D versus 3D SEA *p-values* for each drug-target pair. Tanimoto coefficients score the similarity of 2D versus 3D structures for the searched drug against its most similar known ligand(s) of the target by ECFP (left) and E3FP (right). E3FP uses an early parameter set. Supporting Table S4 shows recalculated SEA *p-values* on the final E3FP parameter set used elsewhere. B-E) Experimentally measured binding curves for tested drugs and reference binders (black) at protein targets B) M_5_, C) α2β4, D) α3β4, and E) α4β4. See Table S7 for more details.

Anpirtoline (CHEMBL1316374) is an agonist of the 5-HT_1B_, 5-HT_1A_, and 5-HT_2_ receptors, and an antagonist of the 5-HT_3_ receptor, with *K* _*i*_’s of 28, 150, 1490, and 30 nM, respectively ^68,69^. However, we predicted it would bind to the nAchRs, of which it selectively bound to α3β4 at a *K*_*i*_ of 3.41 μM and an LE of 0.536 kcal/mol/heavy atom (Figure 4d, Table S7). In this case, the motivating SEA E3FP prediction was for the α4β2 subtype of nAchR, for which the experiment was inconclusive, suggesting either that the ligand reference data from ChEMBL distinguishing these subtypes was insufficient, or that the SEA E3FP method itself did not distinguish among them, and this is a point for further study.

Alphaprodine (CHEMBL1529817), an opioid analgesic used as a local anesthetic in pediatric dentistry ^70^, bound to the muscarinic acetylcholine receptor (mAchR) M_5_ with a *K*_*i*_ of 771 nM and an LE of 0.442 kcal/mol/heavy atom (Figure 4b, Figure S9e). We found no agonist activity on M_5_ by alphaprodine by Tango assay ^71,72^ (Figure S10b), but we did find it to be an antagonist (Figure S11). Intriguingly, alphaprodine also showed no significant affinity for any of the muscarinic receptors M_1_-M_4_ up to 10 μM (Figures S9a-d), indicating that it is an M_5_-selective antagonist. Muscarinic M_5_ selective small molecules are rare in the literature ^73^. Whereas its M_5_ selectivity would need to be considered in the context of its opioid activity (mu, kappa, and delta opioid receptor affinities however are not publicly available), alphaprodine nonetheless may find utility as a M_5_ chemical probe, given the paucity of subtype-selective muscarinic compounds. Interestingly, the E3FP SEA prediction leading us to the discovery of this activity was for the muscarinic M_3_ receptor, to which alphaprodine ultimately did not bind and for which alphaprodine showed no agonist activity (Figure S10a). This highlights not only the limitations of similarity-based methods such as SEA for the discovery of new subtype-selective compounds when none of that type are previously known, but also the opportunity such methods provide to identify chemotypes and overall receptor families that merit further study nonetheless.

## Discussion

Three results emerge from this study. First, we encode a simple three-dimensional molecular representation into a new type of chemical informatic fingerprint, which may be used to compare molecules in a manner analogous to that already used for two-dimensional molecular similarity. Second, the 3D fingerprints contain discriminating information that is naturally absent from 2D fingerprints, such as stereochemistry and relationships among atoms that are close in space but distant in their direct bond connectivity. Finally, as small molecules may adopt many structural conformations, we combine conformation-specific 3D fingerprints into sets to evaluate entire conformational ensembles at once. This may be of interest in cases where different conformations of a molecule are competent at diverse binding sites across the array of proteins for which that same molecule is, at various potencies, a ligand.

We devised a simple representation of three-dimensional molecular structures, an “extended 3D fingerprint” (E3FP), that is directly analogous to gold standard two-dimensional approaches such as the extended connectivity fingerprint (ECFP). As with two-dimensional fingerprints, this approach enables pre-calculation of fingerprints for all conformers of interest in an entire library of molecules once. Unlike conventional 3D approaches, similarity calculations in E3FP do not require an alignment step. Consequently, E3FP similarity calculations are substantially faster than standard 3D comparison approaches such as ROCS. Furthermore, E3FP fingerprints are formatted identically to ECFP and other 2D fingerprints. Thus systems pharmacology approaches such as SEA ^4,5^, Naive Bayes Classifiers ^9^, SVM ^14^, and other established machine learning methods may readily incorporate E3FPs for molecular conformers without modification. While choices of E3FP’s parameter space might be specifically optimized for the machine learning method in question, we have demonstrated that E3FP’s highest-performing parameter choice for SEA (Figure 2c-d) produces fingerprints that likewise perform well for SVM, random forests, and neural networks (Figures S5-S6).

To explore the role of 2D vs 3D features in the discriminatory power of molecular fingerprints, we progressively disabled capabilities specific to E3FP, such as stereochemistry encoding (termed “E3FP-NoStereo”) and non-bonded atom relationships (termed “E2FP-Stereo”), eventually arriving at a stripped-down version of E3FP (termed “E2FP”) that, much like ECFP, encodes only 2D information. We evaluated the consequences of removing these three-dimensional features on performance in retrospective machine learning tasks (e.g., Figure 2c-e; Table 1; Figures S7-S8) We found that inclusion of non-bonded atoms was a more important contributor to performance than stereochemical information. Intriguingly, while progressively adding stereochemical information and inclusion of nonbonded atoms produces marked improvement over ECFP4, inclusion only of nonbonded atom information produces the highest performance fingerprint of all, perhaps because 3D orientations of larger substructures are implicitly encoded within shells purely by relative distances. This observation leads us to believe that a more balanced inclusion of stereochemical information and nonbonded atoms may produce an even higher performing fingerprint. Historically, 3D representations have typically underperformed 2D ones such as ECFP ^7^, and this has always been the case with Similarity Ensemble Approach (SEA) calculations in particular ^6^. Here, however, we find that E3FP exceeds the performance of ECFP in its precision-recall curve (PRC) and matches that of ECFP in its receiver-operating characteristic curve (ROC) area under the curve (AUC) scores (Figure 2c-e; Table 1; Figures S7-S8). While the ROC curve evaluates the general usefulness of the fingerprint for classification by comparing sensitivity and specificity, the precision-recall evaluates how useful the method is for real cases where most tested drug-target pairs are expected to have no affinity. The increased performance in PRC curves when using E3FP over ECFP4 therefore indicates an increased likelihood of predicting novel drug-target pairs that will be experimentally born out with no loss in predictive power.

E3FP’s utility for this task became especially clear when we used it to predict novel drug to protein binding interactions. To do so, we examined only strong SEA predictions with E3FP (SEA-E3FP; *p-value* ≤ 1x10^-25^) that could not be predicted using SEA with ECFP (SEA-ECFP; *p-value* ≥ 0.1). We considered this a challenging task because on-market drugs might be expected to have fewer unreported off-targets in general than a comparatively newer and less-studied research compound might. Furthermore, much of the prior work in chemical informatics guiding molecule design and target testing has been motivated by 2D approaches ^2,7,74^. Accordingly, approximately half of the new predictions were inconclusive in this first prospective test of the method (Tables S4 and S6). Nonetheless, many also succeeded with high ligand efficiencies (LEs), and these included unique selectivity profiles (Figure 4). In one example, SEA-E3FP successfully predicted that alphaprodine would also act as an antagonist of the M_5_ muscarinic receptor, which to our knowledge is not only a new “off-target” activity for this drug, but also constitutes a rare, subtype selective M_5_ antimuscarinic ligand ^73^. The M_5_ muscarinic receptor has roles in cocaine addiction ^75^, morphine addiction ^76^, and dilation of cerebral blood vessels, with potential implications for Alzheimer’s disease ^77^. Study of M_5_ receptors has been hindered by a lack of selective ligands. Due to serious adverse reactions ^78^, alphaprodine was withdrawn from the market in the United States in 1986 and is therefore unlikely to be applied as a therapeutic. However, alphaprodine might be explored not only as a chemical probe for studying M_5_, but also as a reference for future therapeutic development.

Anpirtoline and cypenamine, likewise predicted and subsequently experimentally confirmed to bind previously unreported off-targets among the nicotinic receptors, exhibited exceptional LEs (0.536 - 0.637 kcal/mol/heavy atom), a commonly used metric of optimization potential. Recent patents combining psychostimulants with low-dose antiepileptic agents for the treatment of attention deficit hyperactivity disorder (ADHD) incorporate cypenamine ^79,80^, and nicotinic agents improve cognition and combat ADHD ^81^. Given likewise the association of nicotinic acetylcholine receptor (nAchR) α4 gene polymorphisms with ADHD ^82^, a combination of traditional psychostimulant activity with “non-stimulant” nAChR activity via α4 might improve anti-ADHD efficacy. Whereas cypenamine’s micromolar binding concentration to nAchR is likely below the plasma concentrations it reaches at steady state, its exceptional LEs at nAchR may support further optimization of this pharmacology. As with cypenamine, anpirtoline may serve as a well-characterized starting point for further nAchR optimization, and secondarily, its serotonergic activity may serve as a guide to explore cypenamine’s likely serotonergic activity. Anpirtoline’s benign side effect profile, combined with the nAchR α3β4 subunit’s role in nicotine addiction ^83^ and the lack of α3β4 specific drugs ^84^, motivate further exploration.

We find that, whereas E3FP’s performance matches or exceeds that of ECFP under multiple retrospective metrics, and whereas it leads to new off-target predictions complementing those of ECFP with SEA, there are cases where the more traditional 2D representation yields higher retrospective performance. It would be difficult to tease out the impact that 2D has of necessity made in guiding the design and testing of such molecules, and only time will tell whether ECFP’s higher performance in these cases is due to true pharmacology or historical bias. However, we currently find that ECFP outperforms E3FP on specific targets using SEA (Figure 2f) and in general when applying other machine learning methods (Figures S5-S6). Similarly, ECFP performs well on highly flexible molecules, owing to the difficulty of a small conformer library representing the flexibility of these molecules. Conversely, E3FP’s potential for discerning similar target binding profiles is best realized when comparing molecules with a high degree of conformational similarity on the one hand or on the other one or more chiral centers. As is evident from their respective PRC plots, E3FP typically discriminates SEA predictions more than ECFP does, thereby achieving a better precision-recall ratio, at the initial cost of some sensitivity (Figure 2c). However, this also allows E3FP to consider more distant molecular similarity relationships while maintaining greater discriminatory power than ECFP does at this range. It would be interesting to explore whether some of these more distant relationships might also be regions of pharmacological novelty.

One longtime advantage of 2D molecular representations has been their ability to implicitly sidestep the question of conformation. Whereas heroic effort has gone into solving the crystallographic conformations of hundreds of thousands of small molecules ^85,86^, the binding-competent 3D conformations for millions of research ^25^ and purchasable ^31^ small molecules are not known. Furthermore, polypharmacology exacerbates this problem, wherein a single small molecule can bind many protein partners, as it is not always the case that the molecule in question will adopt the same conformation for each binding site ^2^. Powerful methods to enumerate and energetically score potential conformations exist ^87–89^, but it falls to the researcher to prioritize which of these conformers may be most relevant for a given protein or question. Treating the top five, ten, or more most energetically favorable conformers as a single set, however, may be an alternate solution to this problem. We originally developed SEA so as to compare entire sets of molecular fingerprints against each other ^4^, so it seemed natural to use it in a conformational-set-wise manner here. Furthermore, because SEA capitalizes on nearest-neighbor similarities among ligands across sets of molecules, we expected that it might analogously benefit from nearest-neighbor similarities in conformational space, on a protein-by-protein basis. This may indeed be the case, although we have not attempted to deconvolve E3FP’s performance in a way that would answer whether different E3FPs, and hence different conformations, of the same molecule most account for its predicted binding to different protein targets.

The E3FP approach is not without its limitations. E3FP fingerprints operate on a pre-generated library of molecular conformers. The presence of multiple conformers and therefore multiple fingerprints for a single molecule hampers machine learning performance in naive implementations (Figures S5-S6), as flexible molecules dominate the training and testing data. We anticipate higher numbers of accepted conformers to only exacerbate the problem. The full conformational diversity of large, flexible molecules pose a substantial representational challenge as well (Figure 3c-d). As E3FP depends upon conformer generation, a generator that consistently imposes specific stereochemistry on a center lacking chiral information may produce artificially low or high 3D similarity (Figure 3c). Furthermore, the core intuition of E3FP hinges on the assumption that most binding sites will have differing affinities for molecules with diverging stereochemical orientations, such as stereoisomers. Due to site flexibility, this is not always the case (Figure 3c-d).

Despite these caveats, we hope that this simple, rapid, and conformer-specific extended three-dimensional fingerprint (E3FP) will be immediately useful to the broader community. To this end, we have designed E3FP to integrate directly into the most commonly used protein target prediction methods without modification. An open-source repository implementing these fingerprints and the code to generate the conformers used in this work is available at <u>https://github.com/keiserlab/e3fp/tree/1.0</u>.

## Experimental Section

### Generating Conformer Libraries

To maximize reproducibility, we generated conformers following a previously published protocol ^35^ using RDKit ^90^. For each molecule, the number of rotatable bonds determined the target number of conformers, *N*, such that: *N*=50 for molecules with less than 8 rotatable bonds, *N*=200 for molecules with 8 to 12 rotatable bonds, and *N*=300 for molecules with over 12 rotatable bonds. We generated a size *2N* pool of potential conformers.

After minimizing conformers with the Universal Force Field ^89^ in RDKit, we sorted them by predicted energy. The lowest energy conformer became the seed for the set of accepted conformers. We considered each candidate conformer in sorted order, calculated its root mean square deviation (RMSD) to the closest accepted conformer, and added the candidate to the accepted set if its RMSD was beyond a predefined distance cutoff *R*. Optionally, we also enforced a maximum energy difference *E* between the lowest and highest energy accepted conformers. After having considered all *2N* conformers, or having accepted *N* conformers, the process terminated, yielding a final set of conformers for that molecule.

We tuned this protocol using three adjustable parameters: (1) the minimum root mean square distance (RMSD) between any two accepted conformers, (2) the maximum computed energy difference between the lowest energy and highest energy accepted conformers, and (3) the number of lowest energy conformers to be accepted (fingerprinted). We generated two different conformer libraries by this protocol. In the first (rms0.5), we used a RMSD cutoff *R*=0.5, with no maximum energy difference *E*. In the second (rms1_e20), we chose a RMSD cutoff *R*=1.0, with a maximum energy difference of 20 kcal/mol.

### Enumerating Protonation States

Where specified, we generated dominant tautomers at pH 7.4 from input SMILES using the CXCALC program distributed with ChemAxon’s Calculator Plugins ^91^. We kept the first two protonation states with at least 20% predicted occupancy. Where no states garnered at least 20% of the molecules, or where protonation failed, we kept the input SMILES unchanged. Conformer generation for each tautomer proceeded independently and in parallel.

### ECFP Fingerprinting

To approximate ECFP fingerprints, we employed the Morgan fingerprint from RDKit using default settings and an appropriate radius. ECFP4 fingerprints, for example, used a Morgan fingerprint of radius 2. Where ECFP with stereochemical information is specified, the same fingerprinting approach was used with chirality information incorporated into the fingerprint.

### E3FP Fingerprinting

Given a specific conformer for a molecule, E3FP generates a 3D fingerprint, parameterized by a shell radius multiplier *r* and a maximum number of iterations (or level) *L*, analogous to half of the diameter in ECFP. E3FP explicitly encodes stereochemistry.

### Generating Initial Identifiers

Like ECFP, E3FP generation is an iterative process and can be terminated at any iteration or upon convergence. At iteration 0, E3FP generation begins by determining initial identifiers for each atom based on six atomic properties, identical to the invariants described in ^32^ : the number of heavy atom immediate neighbors, the valence minus the number of neighboring hydrogens, the atomic number, the atomic mass, the atomic charge, the number of bound hydrogens, and whether the atom is in a ring. For each atom, the array of these values are hashed into a 32-bit integer, the atom identifier at iteration 0. While the hashing function is a matter of choice, so long as it is uniform and random, this implementation used MurmurHash3 ^92^.

### Generating Atom Identifiers at Each Iteration

At each iteration *i* where *i* > 0, we consider each atom independently. Given a center atom, the set of all atoms within a spherical shell of radius *i · r* centered on the atom defines its immediate neighborhood, where the parameter *r* is the shell radius multiplier (Figure 1a). We initialize an array of integer tuples with a number pair consisting of the iteration number *i* and the identifier of the central atom from the previous iteration.

For each non-central atom within the shell, we add to the array an integer 2-tuple consisting of a connectivity identifier and the atom's identifier from the previous iteration. The connectivity identifiers are enumerated as an expanded form of those used for ECFP: the bond order for bond orders of 1-3, 4 for aromatic bonds, and 0 for neighbors not bound to the central atom. To avoid dependence on the order in which atom tuples are added to the array, we sort the positions of all but the first tuple in ascending order. 3-tuples are then formed through the addition of a stereochemical identifier, followed by re-sorting. This process is described in detail below.

We then flatten the completed array into a one-dimensional integer array. We hash this 1D array into a single new 32-bit identifier for the atom and add it to an identifier list for the iteration, after optional filtering described below.

### Adding Stereochemical Identifiers

We generate stereochemical identifiers by defining unique axes from the sorted integer 2-tuples from the previous step combined with spatial information. First, we determine the vectors from the center atom to each atom within the shell. Then, we select the first unique atom by atom identifier from the previous iteration, if possible, and set the vector from the central atom to it as the *y*-axis. Where this is not possible, we set the *y*-axis to the average unit vector of all neighbors. Using the angles between each unit vector and the *y*-axis, the atom closest to 90 degrees from the *y*-axis with a unique atom identifier from the previous iteration defines the vector of the *x*-axis (Figure 1a).

We then assign integer stereochemical identifiers *s*. Atoms in the *y* > 0 and *y* < 0 hemispheres have positive and negative identifiers, respectively. *s*=±1 is assigned to atoms whose unit vectors fall within 5 degrees of the *y*-axis. We divide the remaining surface of the unit sphere into eight octants, four per hemisphere. The *x*-axis falls in the middle of the *s*=±2 octants, and identifiers ±3-5 denote remaining octants radially around the *y*-axis (Figure 1a). If unique *y*- and *x*-axes assignment fails, all stereochemical identifiers are set to 0.

Combining the connectivity indicator and atom identifier with the stereochemical identifier forms a 3-tuple for each atom, which, when hashed, produces an atom identifier dependent orientation of atoms within the shell.

### Removing Duplicate Substructures

Each shell has a corresponding *substructure* defined as the set of atoms whose information is contained within the atoms in a shell. It includes all atoms within the shell on the current iteration as well as the atoms within their substructures in the previous iteration. Two shells have the same substructure when these atom sets are identical, even when the shell atoms are not. As duplicate substructures provide little new information, we filter them by only adding the identifiers to that iteration’s list that correspond to new substructures or, if two new identifiers correspond to the same substructure, the lowest identifier.

### Representing the Fingerprint

After E3FP runs for a specified number of iterations, the result is an array of 32-bit identifiers. We interpret these as the only “on” bits in a *2*^32^length sparse bitvector, and they correspond to 3D substructures. As with ECFP, we “fold” this bitvector to a much smaller length such as 1024 by successively splitting it in half and conducting bitwise OR operations on the halves. The sparseness of the bitvector results in a relatively low collision rate upon folding.

### Fingerprint Set Comparison with SEA

The similarity ensemble approach (SEA) is a method for searching one set of bitvector fingerprints against another set ^4^. SEA outputs the maximum Tanimoto coefficient (TC) between any two fingerprint sets and a *p-value* indicating overall similarity between the sets. SEA first computes all pairwise TCs between the two fingerprint sets. The sum of all TCs above a preset pairwise TC threshold *T* defines a *raw score*. For a given fingerprint, SEA calculates a background distribution of raw scores empirically ^4^. This yields an observed *z-score* distribution, which at suitable values of *T* follows an extreme value distribution (EVD). For values of *T* ranging from 0 to 1, comparing goodness of fit (*chi-square*) to an EVD vs a normal distribution determines an optimal range of *T*, where the empirical *z-score* distribution favors an EVD over a normal distribution. In this EVD regime we may convert a *z-score* to a *p-value* for any given set-set comparison.

### K-fold Cross-Validation with SEA

We performed *k*-fold cross-validation on a target basis by dividing the ligands of at least 10 μM affinity to each target into *k* sets per target. For a given fold, *k-1* ligand sets and their target labels together formed the training data. The remaining ligand sets and their target labels formed the test data set. Due to the high number of negative examples in the test set, this set was reduced by ∼25% by removing all negative target-molecule pairs that were not positive to any target in the test set. Conformers of the same ligand did not span the train vs test set divide for a target. For each fold, conformer fingerprint sets for molecules specific to the test set were searched against the union of all training conformer fingerprints for that target, yielding a molecule-to-target SEA *p-value*. From the -log*p-values* for all test-molecule-to-potential-target tuples, we constructed a receiving operator characteristic (ROC) curve for each target, and calculated its area under the curve (AUC). We likewise calculated the AUC for the Precision-Recall Curve (PRC) at each target. For a given fold, we constructed an ROC curve and a PRC curve using the -log *p-values* and true hit/false hit labels for all individual target test sets, which we then used to compute a fold AUROC and AUPRC. We then computed an average AUROC and AUPRC across all *k* folds. The objective function AUC_SUM_ consisted of the sum of the average AUROC and AUPRC.

### Optimizing Parameters with Spearmint

E3FP fingerprints have the following tunable parameters: stereochemical mode (on/off), nonbound atoms excluded, shell radius multiplier, iteration number, and folding level. Additional tunable parameters for the process of conformer generation itself are the minimum RMSD between conformers, the maximum energy difference between conformers, and how many of the first conformers to use for searching. This parameter space forms a 8-dimensional hypercube. Of the 8 dimensions possible, we employed the Bayesian optimization program Spearmint ^45^ to explore four: shell radius multiplier, iteration number, number of first conformers, and two combinations of values for the RMSD cutoff and maximum energy difference between conformers. We evaluated the parameter sets by an objective function summing ROC and PRC AUCs (AUC_SUM_), and Spearmint proposed future parameter combinations. The objective function evaluated *k*-fold cross-validation with the similarity ensemble approach (SEA) as described in the following section.

For the first stage, the dataset consisted of 10,000 ligands randomly chosen from ChEMBL17, the subset of targets that bound to at least 50 of these ligands at 10 μM or better, and the objective function used was the AUPRC. Spearmint explored values of the shell radius multiplier between 0.1 and 4.0 Å, the number of lowest energy conformers ranging from 1 to all, and maximum iteration number of 5. Additionally, two independent conformer libraries were explored: rms0.5 and rms1_e20 (see above). 343 unique parameter sets were explored. We found that the best parameter sets used less than 35 of the lowest energy conformers, a shell radius multiplier between 1.3 and 2.8 Å, and 2-5 iterations. The conformer library used did not have an apparent effect on performance (data not shown).

For the second stage, we ran two independent Spearmint trajectories with a larger dataset consisting of 100,000 ligands randomly chosen from ChEMBL20, the subset of targets that bound to at least 50 of these ligands at 10 μM or better, and the AUC_SUM_ objective function. We employed the CXCALC program ^91^ to determine the two dominant protonation states for each molecule at physiological pH, and then conformers were generated using an RMSD cutoff of 0.5. The number of fingerprinting iterations used in both trajectories was optimized from 2 to 5, but the two trajectories explored different subsets of the remaining optimal parameter ranges identified during the first stage: the first explored shell radius multipliers between 1.3 and 2.8 Å with number of conformers bounded at 35, while the second explored shell radius multipliers between 1.7 and 2.8 Å with number of conformers bounded at 20. Spearmint tested 100 parameter combinations in each trajectory.

During optimization, we observed that the simple heuristic used by SEA to automatically select the TC threshold for significance resulted in folds with high TC cutoffs having very high AUPRCs but low AUROCs due to low recall, while folds with low TC cutoffs had lower AUPRCs but very high AUROCs (Figure S3). Several folds in the latter region outperformed ECFP4 in both AUPRC and AUROC (Figure S3c). We therefore selected the best parameter set as that which produced the highest AUC_SUM_ while simultaneously outperforming ECFP4 in both metrics. For all future comparisons, the TC cutoff that produced the best fold results was applied to all folds during cross-validation.

### K-fold Cross-Validation with Other Classifiers

We performed k-fold cross-validation using alternative classifiers in the same manner as for SEA, with the following differences. We trained individual classifiers on a target by target basis. In the training and test data, we naively treated each conformer fingerprint as a distinct molecular fingerprint, such that the conformer fingerprints did not form a coherent set. After evaluating the target classifier on each fingerprint for a molecule, we set the molecule score to be the maximum score of all of its conformer fingerprints.

For the Naive Bayes (NB), Random Forest (RF), and Support Vector Machine with a linear kernel (LinSVM) classifiers, we used Scikit-learn version 0.18.1 (<u>https://github.com/scikit</u>-learn/scikit-learn/tree/0.18.1). We used default initialization parameters, except where otherwise specified. For the RF classifier, we used 100 trees with a maximum depth of 2. We weighted classes (positive and negative target/molecule pairs) to account for class imbalance. For LinSVM kernel, we applied an l1 norm penalty and balanced class weights as for RF.

We implemented Artificial Neural Network (NN) classifiers with nolearn version 0.6.0 (<u>https://github.com/dnouri/nolearn/tree/0.6.0</u>). We trained networks independently for each target using 1024-bit input representations from either E3FP or ECFP. The NN architecture comprised 3 layers: an input layer, a single hidden layer with 512 nodes, and an output layer. We used dropout ^93^ as a regularizer on the input and hidden layers at rates of 10% and 25%, respectively. The hidden layer activation function was Leaky Rectified Linear ^94^ with default leakiness of 0.01. The prediction layer used softmax nonlinearities. We trained networks trained for 1000 epochs with early stopping to avoid overfitting, by monitoring the previous 75 epochs for lack of change in the loss function. The final softmax layer contained 2 tasks (classes), one corresponding to binding and the other corresponding to the absence of binding. This softmax layer produced a vector corresponding to the probability of a given molecule being a binder or non-binder given the neural network model. We calculated training error using a categorical cross entropy loss.

### Predicting Novel Compound-Target Binding Pairs

To identify novel compound-target pairs predicted by E3FP but not by ECFP, we built a subset of 309 proteins/complex mammalian targets (106 human) for which the National Institute of Mental Health Psychoactive Drug Screening Program (NIMH PDSP) ^63^ had established binding assays. We selected all compounds listed as in-stock in ZINC15 ^31^, downloaded on 2015-09-24. We fingerprinted all ligands in ChEMBL20 ^95^ with affinity < 10 μM to the PDSP targets using the RDKit Morgan algorithm (an ECFP implementation) as well as by a preliminary version of E3FP (Table S4). We likewise fingerprinted the ZINC15 compounds using both ECFP and E3FP. We queried the search compounds using SEA against a discrete sets of ligands from < 10 nM affinity (strong binders) to < 10 μM affinity (weak binders) to each target, in log-order bins, using both ECFP and E3FP independently. We filtered the resulting predictions down to those with a strong SEA-E3FP *p-value* (< 1x10^-25^) and ≤ 10 nM affinity to the target, where the SEA-ECFP *p-value* exceeded 0.1 (i.e., there was no significant SEA-ECFP prediction) in the same log-order affinity bin. From this set of compound-target pairs, we manually selected eight for experimental testing.

### Experimental Assays of Compound-Target Binding Pairs

Radioligand binding and functional assays were performed as previously described ^71,96,97^. Detailed experimental protocols and curve fitting procedures are available on the NIMH PDSP website at: <u>https://pdspdb.unc.edu/pdspWeb/content/PDSP%20Protocols%20II%202013-03-28.pdf</u>.

Ligand efficiencies were calculated using the expression

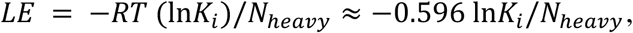

where *R* is the ideal gas constant, *T* is the experimental temperature in Kelvin, and *N*_*heavy*_ is the number of heavy atoms in the molecule ^98^. The ligand efficiency is expressed in units of kcal/mol/heavy atom.

### Source Code

Code for generating E3FP fingerprints is available at <u>https://github.com/keiserlab/e3fp/tree/1.0</u> under the GNU Lesser General Public License version 3.0 (LGPLv3) license. All code necessary to reproduce this work is available at <u>https://github.com/keiserlab/e3fp-paper/tree/1.0</u> under the GNU LGPLv3 license.

## Supporting Information

Supporting figures and tables include an enlarged Figure 1c, parameter optimization and cross-validation results, references for highlighted molecule pairs in Figure 3, descriptions of compounds used in experiments, and all experimental results. This material is available free of charge via the Internet at http://pubs.acs.org.

## Acknowledgments

This material is based upon work supported by a Paul G. Allen Family Foundation Distinguished Investigator Award (to MJK), a New Frontier Research Award from the Program for Breakthrough Biomedical Research, which is partially funded by the Sandler Foundation (to MJK), and the National Science Foundation Graduate Research Fellowship Program under Grant No. 1650113 (to SDA and ELC). ELC is a Howard Hughes Medical Institute Gilliam Fellow. *K*_*i*_ determinations and agonist and antagonist functional data was generously provided by the National Institute of Mental Health's Psychoactive Drug Screening Program, Contract # HHSN-271-2013-00017-C (NIMH PDSP). The NIMH PDSP is Directed by Bryan L. Roth MD, PhD at the University of North Carolina at Chapel Hill and Project Officer Jamie Driscoll at NIMH, Bethesda MD, USA. We thank Teague Stirling, Michael Mysinger, Cristina Melero, John Irwin, William DeGrado, and Brian Shoichet for discussions and technical support.

## Abbreviations Used

AUPRC: AUC of the Precision-Recall Curve
AUROC: AUC of the Receiver Operating Characteristic Curve
E3FP: Extended Three-Dimensional FingerPrint
ECFP: Extended Connectivity FingerPrint
NB: Naive Bayes Classifier
NN: Artificial Neural Network
PRC: Precision-Recall Curve
RF: Random Forest
ROC: Receiver Operating Characteristic Curve
SEA: Similarity Ensemble Approach
SVM: Support Vector Machine
TC: Tanimoto coefficient

